# Nanovibrational stimulation of osteogenesis engages non-canonical Wnt signalling and NF-κB regulator BCL3 as a mechanotransducer

**DOI:** 10.1101/2025.07.24.666351

**Authors:** Hussain Jaffery, Udesh Dhawan, Jonathan A Williams, Carmen Huesa, Ruaidhrí J Carmody, James FC Windmill, Stuart Reid, Peter Childs, Manuel Salmeron-Sanchez, Matthew J Dalby

## Abstract

Enhancing osteogenesis in mesenchymal stromal (stem) cells is essential for advancing cellular therapies that target and alleviate skeletal pathologies. By employing a bespoke bioreactor capable of delivering nano-amplitude vibration (30 nm, 1kHz) to human adipose-derived mesenchymal stromal cells, we observed osteo-specific differentiation. We related this to the mechanotransductive mechanism by showing that inhibition of intracellular tension results in the loss of cytoskeletal organisation and myosin activation driven by nanovibration. Further, we dissect the mechanism of osteogenesis using a panel of Wnt agonists and antagonists and highlight the role of non-canonical Wnt. Then, using *Bcl3*^-/-^ cells and by stimulating with BCL3 peptide, we show that non-canonical Wnt, osteogenesis-related inflammation and osteogenesis itself are all regulated by BCL3. This is important as nanoscale direct cell-stimulation is gaining interest, and there is an emerging consensus that such signals can be osteogenic. While prior research has only broadly hinted at how nanovibrational signals convert to an osteogenic phenotype, this new work pinpoints critical mechanistic insights, thereby advancing our understanding of this promising avenue in musculoskeletal cell therapy.

## Introduction

Pathologies of the skeletal system, comprising bone and joint diseases, affect at least 1.7 billion people worldwide.^1^ Low bone density and reduced bone formation afflicts a large subset of patients with congenital and emergent diseases, such as osteogenesis imperfecta and osteoporosis, in elective procedures such as with reduced bone stock around joint replacement implants, and in traumatic injuries resultant in non-union fractures.^2,3^ Provision of cellular therapies, typically focusing on mesenchymal stromal (or stem) cells (MSCs) that can differentiate into bone forming osteoblasts is an emerging focus.^4,5^ For cell therapy, cell supply is key and so there is considerable focus on MSCs derived from abundant reservoirs in adipose tissue, multipotent adipose-derived stromal cells (ADSCs).^4,6^ The MSC/ADSC approach allows for both autologous and allogeneic therapy approaches due to the immune modifying effects of these stromal cells.^7,8^

Typically, osteogenic differentiation of MSCs/ADSCs is driven using soluble factors such as corticosteroids (e.g. dexamethasone) or growth factors (e.g. bone morphogenetic protein 2, BMP-2.^6,9^ In cell therapy manufacture, these approaches can be problematic as biologics that have not previously been used in GMP (good manufacturing practice) production of therapeutic MSC/ADSC derived cells need to be sourced and regulated. Further, these factors can non-specifically activate disparate cell programmes with lineage ambiguity and deleterious effects such as the promotion of cancer.^9^ We, therefore, advocate a biomechanical approach to MSC/ADSC differentiation that uses only mechanical displacement at the nanometre scale. Application of low amplitude vibration and nanovibrational (NV) signals, has been shown to be osteogenic and has the advantage that therapies can be manufactured in flasks and using media that have been previously regulated in the production of human therapies.^4,10–13^

We have designed a bespoke bioreactor device that provides a piezoelectrically-driven oscillating displacement of 30 nm at 1 kHz in the vertical z-axis to any affixed cell culture plasticware, with a force of 5 nN.^14^ Our NV approach has further benefits compared to use of soluble factors with highly targeted osteogenic differentiation, e.g., in reducing co-stimulation of adipogenesis and limiting off-target differentiation.^15,16^ However, the underlying causal molecular mechanisms have remained elusive.

Emerging literature has previously shown that nano-vibrational (NV) stimulation strengthens focal-adhesion complexes and remodels the actin cytoskeleton, funnelling these mechanical cues into the mitogen-activated protein-kinase (MAPK) network including the ERK1/2 cascade. Nuclear ERK1/2 then phosphorylates and potentiates RUNX2, the master osteogenic transcription factor, thereby switching on bone-specific gene programs and accelerating osteogenesis.^15,17^ Yes-associated protein (YAP) translocation to the nucleus has also been linked to changes in adhesion and cytoskeleton and activation of RUNX2.^18^ Further, adhesion-cytoskeleton linked to mechano-regulated ion channels, such as PIEZO and transient receptor potential channels, have been implicated with osteogenesis through activation of canonical Wnt (portmanteau of wingless and Int-1) - β-catenin signalling, activating RUNX2.^15^

In this study, we seek to dissect molecular control of nanovibrationally-directed osteogenesis with a particular focus on canonical and non-canonical Wnt signalling. We establish that precise nanovibration promotes cytoskeletal reorganisation and enhances expression of an axis of proinflammatory, non-canonical Wnt and osteogenic genes expression, and further show that the NF-κB regulating protein BCL3 is pivotal to coordinating this gene expression signature.

## Results and Discussion

### High-precision nanovibration generation by bioreactors and nanoforce delivery to cell cultureware

A nanovibrational bioreactor was used to deliver vibration to ADSCs (Figure 1a).^14^ The bioreactor was designed to deliver a consistent and precise vertical pistonic vibration, with an amplitude of 30 nm and frequency of 1 kHz, to cells adhered to standard cell culture plates. The term pistonic is used to indicate this piston-like motion of the top plate of the bioreactor which vibrates at 1 kHz with the same amplitude across its surface.

**Figure 1.**
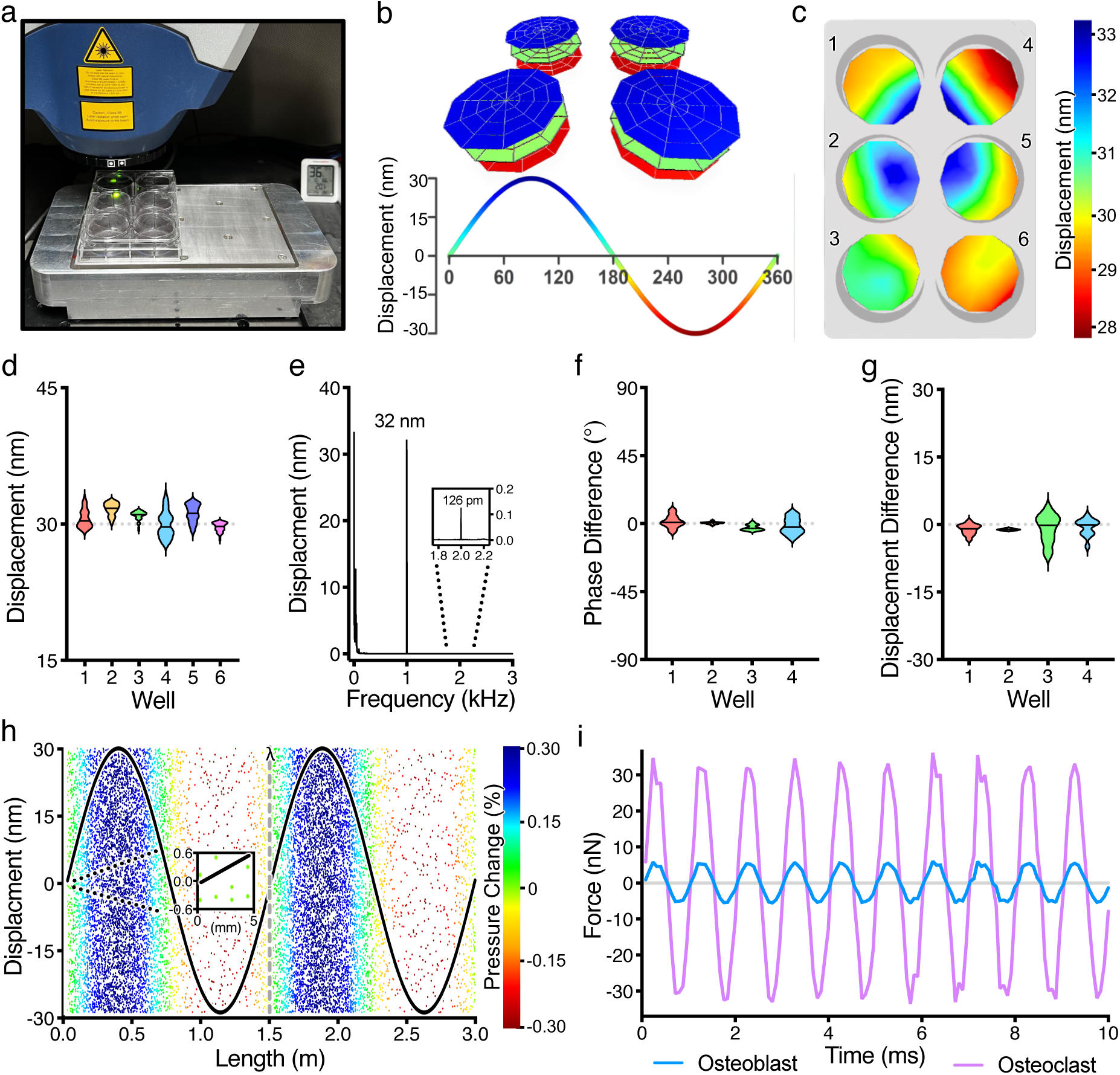
Scanning laser Doppler vibrometry-based characterisation of the nanovibrational bioreactor. The nanovibrational bioreactor and 6-well plate magnetically coupled to the top plate (a). The instantaneous z-displacement at three separate points (0 (green), 90° (blue) and 270° (red) of one vibration cycle, measured at 1 kHz (b). Colour map of maximum z-displacement (90° of vibration cycle) in relation to position within wells of culture plate at 1 kHz (c). Violin plot of the displacement at 1 kHz for each well of the 6-well plate, with 41 measurement sites per well (d). The average frequency spectrum of the z-displacement from 0.4 to 3000 Hz from all measurement points, with a primary peak at 1 kHz and a much lower 2 kHz harmonic (e). The phase difference (f) and displacement difference (g) between the bottom of each well and the cell media’s surface. The acoustic wave pressure distribution in cell media across two wavelengths is plotted as a point cloud representation, where point density and colour represent the spatial variability, with vibration displacement superimposed on top (h). The zoomed-in section (inset with dotted lines) depicts the very minimal pressure change across the 5 mm of cell media used within each well and the labelled vertical grey dashed line indicates one wavelength (λ). The estimated accelerative force experienced by individual stromal cells/osteoblasts and osteoclasts adhered to the bottom of a 6-well plate, based on the mass of cell media above each cell-type on average (i).

We utilised scanning laser Doppler vibrometry to precisely characterise the combined setups with the bioreactor: a 6-well plate (Figure 1) or a 4-well chamber slide (supplemental Figure 1) used in this work. The pistonic nature of the vibration is shown in Figure 1b, which depicts the instantaneous displacement across 4 wells of the 6-well plate atop the nanovibrational bioreactor at three separate points of the vibration cycle (0°, 90°and 270°). For a 6-well plate mounted to the nanovibrational bioreactor, a maximum 5 nm relative variation within individual wells was observed, (Figure 1c) and a 2 nm variation on average between the wells (Figure 1d) in maximum vertical displacement at 1 kHz was observed.

In the average frequency spectrum of the combined setup, we observe two regions: a low-frequency section caused by seismic/ambient background noise and the desired ∼30 nm nanovibration peak at 1 kHz, representing a pure audible tone (Figure 1e). The bioreactor produces minimal detectable harmonics, as evidenced by the ∼250X lower ∼120 picometer displacement harmonic at 2 kHz, which is hard to distinguish from the background, indicating that the bioreactor only mechanically stimulates cells at 1 kHz (Figure 1e).

There is no displacement or phase difference (*P* > 0.05) between the bottom of the well and the cell media’s surface (Figures 1f and 1g), indicating that the cell media vibrates as a rigid body (i.e., incompressibly) in response to the applied nanoamplitude vibration, with its surface vibrating in unison with the bottom of the culture ware. This suggests that viscous and shear forces are negligible, a consequence of the wavelength of the acoustic nanoamplitude vibration (∼1.5 m in cell media) being orders of magnitude larger than the height of cell media of ∼5 mm (Figure 1h), indicating a very minimal change in instantaneous acoustic pressure within the media. The maximum pressure change caused by nanovibration is 283 Pa, which equates to a ±0.3% change in ambient pressure. While the maximum pressure change over the 5 mm of cell media is even more negligible (6 Pa). This enabled us to calculate using Newton’s second law, that the nanovibration (1 kHz, ±30 nm) exerts a nanoscale sinusoidal accelerative force of approximately ±5 nN per stromal/osteoblast cell and ±32 nN per average osteoclast cell (Figure 1i). The 4-well chamber slide provides a similar level of displacement accuracy at 1 kHz (see supplemental Figure 1). Thus, our high-precision nanovibrational bioreactor serves as a platform to probe cellular mechanotransduction with nanonewton-scale forces.

### Nanovibrational stimulation elicits ADSC cytoskeletal reorganisation and mechanotransductive responses

To characterise the biomechanical response of cells to the nanovibrational stimulus, immunofluorescent assessment of the cytoskeleton was undertaken (Figure 2). The actin cytoskeleton directly organises to support external mechanical force and also dynamically responds via latent biochemical modifications, making it the ideal cellular organelle to assess force-response.^19^ The nanoforce delivered in the nanovibrational stimulation of ADSCs enhanced expression and colocalisation of phosphorylated (Ser19) myosin light chain-2 (p-MLC2) with filamentous actin (F-Actin) bundles, an association that organises cell cytoskeleton via activation of mechanosensory Rho-associate protein kinase (ROCK) signalling (Figure 2a,b). Inhibition of ROCK activity with Y27632, in the presence of nanovibration, reversed the co-localisation of F-Actin and p-MLC2 and reduced their expression levels (Figure 2c). Immunoblot analysis confirmed that ROCK inhibition substantially reversed the nanovibration-driven enhancement of protein expression levels (Figure 2d and supplemental Figure 2). Functionally, p-MLC2 is downstream to ROCK and controls actomyosin contraction, while cell division control protein 42 homolog (CDC42) is a Rho GTPase and is downstream to PKC in the Wnt-Ca^2+^ pathway.^20,21^ Phosphorylated Cofilin (p-Cofilin) is itself downstream to CDC42 and maintains actin polymerisation.^20,21^ The increase in these cytoskeletal markers during nanovibrational stimulation and reversal with ROCK (downstream in the non-canonical Wnt-planar cell polarity, PCP, pathway) inhibition (Figure 2d) is strongly indicative of the involvement of non-canonical Wnt signalling being central to nanovibrational osteogenesis, which we investigated next.^21^

**Figure 2.**
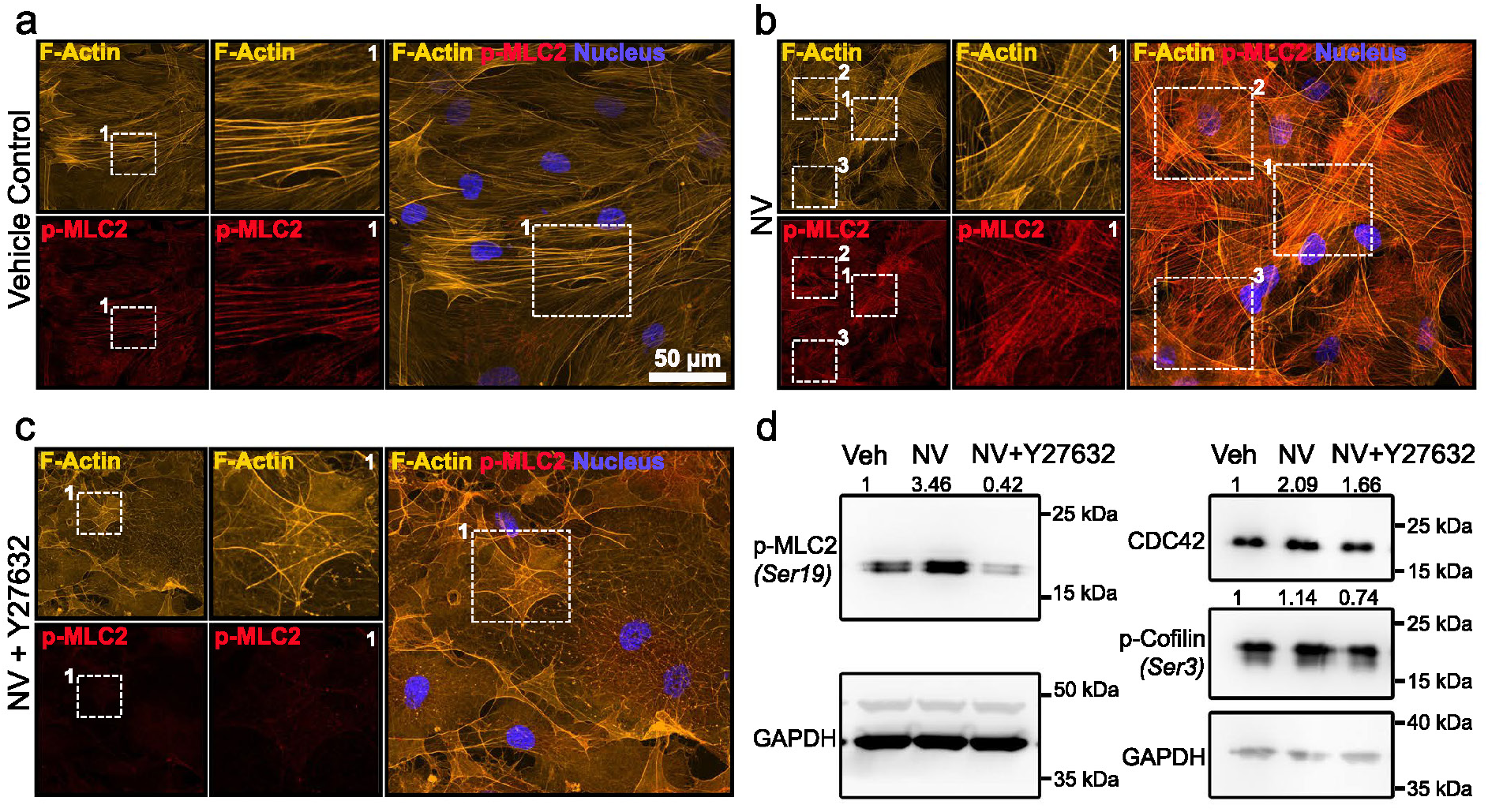
Rho-associated coiled-coil kinases (ROCK) are mechanosensors of nanovibrational stimulation. Immunofluorescent images showing baseline expression of cytoskeletal proteins filamentous actin (F-Actin) and phosphorylated myosin light chain-2 (pMLC2) in vehicle control (H_2_O) adipose-derived stem cells (ADSCs) (a). Immunofluorescent images showing enhanced colocalization of F-Actin and p-MLC2 upon nanovibrational (NV) stimulation, confirming that nanovibration activates ROCK signalling (b). Immunofluorescent images showing F-Actin disruption and reduced p-MLC2 expression upon treatment with ROCK inhibitor, Y27632 (c). All images are representative of observations from three independent biological replicates. Immunoblots confirming that nanovibrational stimulation regulates cytoskeleton-regulating protein CDC42 and activates ROCK mechanosensing by phosphorylating MLC2 and Cofilin (d). Numerical values above cropped, representative immunoblots are quantitative averages from n = 3 independent biological replicates. The uncropped immunoblots are shown in supporting information.

### Nanovibrational stimulation induces ADSC osteogenesis through expression of proinflammatory, Wnt-pathway and osteogenic genes

To assess the ability and efficacy of nanovibration in inducing osteogenesis, we compared it to the benchmark metabolite media (MM, or osteogenic media) based osteogenic induction (Figure 3). Utilised for mammalian osteoblast *in vitro* differentiation, MM incorporates ascorbic acid, β-glycerophosphate and dexamethasone, each with multiple mechanistic targets.^22–24^

**Figure 3.**
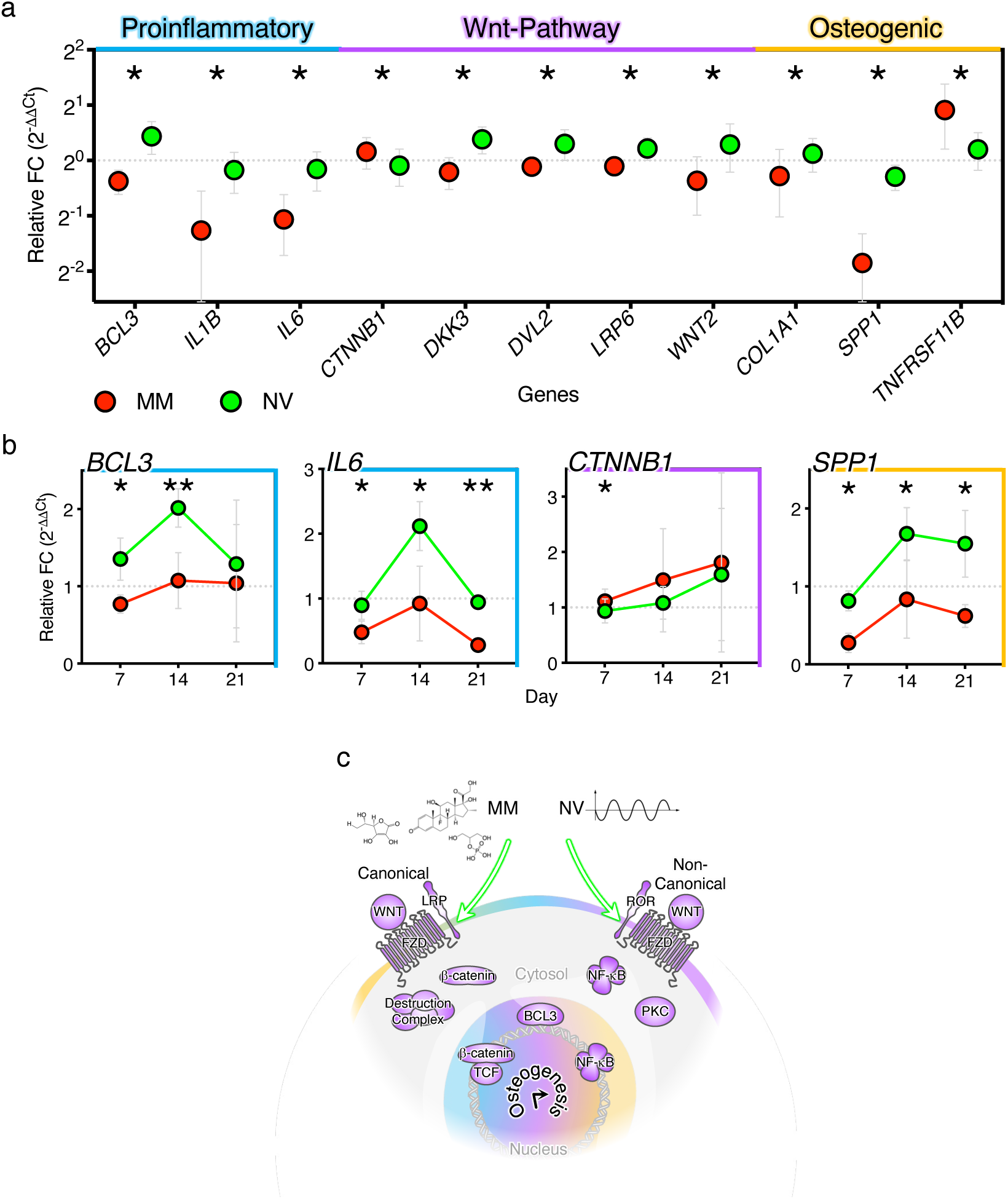
Nanovibration elicits a unique gene expression signature involving proinflammatory, Wnt-pathway and osteogenic markers. Human adipose-derived stromal cells (ADSCs) that were stimulated with nanovibration (NV) compared to osteogenic metabolite medium (MM), at day 7 (a), 14 or 21 (b) were assessed for transcriptional gene mRNA expression via relative quantitative-PCR. Both groups were normalised to gene expression of basal unstimulated cells (baseline at 1). Repeat-measure one-way analysis of variance with Fisher’s least significant difference test incorporating basal unstimulated group (a) or paired t-tests (b) of negative delta-delta Ct values, n = 3 independent biological replicates. Schematic representation summarising the impact of nanovibration increasing expression of gene sets categorised as proinflammatory, of the Wnt pathway or osteogenic markers, while metabolite medium has relatively decreased expression (c).

First, we considered inflammation. The proinflammatory NF-κB (nuclear factor kappa-light-chain enhancer of activated B cells) transcription factor family act as a regulator of cell differentiation decisions and regulates expression of proinflammatory cytokines such as interleukin (IL)-1β or IL-6.^25^ BCL3 (B-cell lymphoma 3-encoded protein 3) acts as a transcriptional co-regulator of NF-κB.^26,27^ Compared to MM osteogenic induction, cells stimulated with nanovibration for 7 days expressed significant increases in NF-κB related proinflammatory genes *BCL3*, *IL1B* and *IL6* (Figure 3a, maximal expression of BCL3 and IL6 was seen at day 14, Figure 3b). Active cell processes, such as differentiation stimuli, wound healing and bone repair employ low, controlled levels of stromal cell inflammatory cytokine expression, contextualising the proinflammatory gene activation seen in these naïve ADSCs.^16,28–31^

Developmental Wnt (Wingless and Int-1) ligand signalling and its associated pathway plays an integral role in spatiotemporal gene expression and cell differentiation.^32^ Wnt ligands bind to frizzled (Fz) G protein-coupled receptors, canonical Wnt involves β-catenin transcription factor activity and its restriction by the β-catenin destruction complex, whereas non-canonical Wnt incorporates the Ca^2+^-NFAT (nuclear factor of activated T-cells) signalling pathway branch and the Rho (Ras Homolog Family)/Rac (Rac Family Small GTPase) transducers and AP-1 (activator protein 1) transcription factor, in the planar cell polarity (PCP) pathway branch.^32^ Here we see repression of β-catenin (*CTNNB1*) for nanovibrational stimulation compared to MM stimulation of ADSCs, indicating canonical signalling is repressed in nanovibrational stimulation (Figure 3a at day 7 and Figure 3b showing this repression is maintained with time in culture). As our hypothesis emerging from Figure 3 is that non-canonical Wnt is important, this is perhaps sensible. This hypothesis is further supported as transcripts for *DKK3* (Dickkopf-related protein 3), *DVL2* (segment polarity protein dishevelled homolog), *LRP6* (low-density lipoprotein receptor-related protein 6) and *WNT2* (wingless-type MMTV integration site family, member 2) were all up-regulated by nanovibrational stimulation and are all linked to non-canonical Wnt signalling.^33,34^

Nanovibrational stimulation of MSCs has been linked in several reports to turn-on the osteogenic programme, and again, here transcripts related to osteogenesis (collagen I, *COL1A1*, the major bone ECM protein) and osteopontin (secreted phosphoprotein 1, *SPP1*, a bone ECM protein) were seen to be upregulated with nanovibration compared to MM treated control cells at day 7 (Figure 3a,b), with maximal expression at day 14 for each that was tested at 3 time points (Figure 3b). Osteoprotegerin (encoded by the tumour necrosis factor receptor superfamily member 11B, gene, *TNFRSF11B*) that is expressed by bone forming cells to reduce bone removing osteoclastic cell activity,^35^ was seen to have reduced expression at day 7 in nanovibrational stimulated cells compared to MM treated control cells (Figure 3a). Gene transcripts tested but not significantly changed at day 7 are shown in supplemental Figure 3). Taken together, these data show proinflammatory, Wnt, and osteogenic gene expression is emergent in early differentiation through nanovibrational stimulation, compared to MM induced differentiation (Figure 3c).

### Inhibition of Wnt implicates the non-canonical Wnt pathway in driving nanovibrational osteogenesis

Having seen that nanovibration triggers a distinct proinflammatory and Wnt-related gene signature compared to chemical osteogenic media, we next wanted to elucidate which branch of the Wnt signalling pathway was activated in nanovibrational stimulation during very early differentiation. ADSCs were treated with small-molecule inhibitors targeting specific branches of Wnt pathway regulation and function, in the presence of nanovibration, for 24 hours (Figure 4). Inhibitor LGK974 (LGK) restricts pan-Wnt extracellular secretion and autocrine/paracrine signalling by targeting and preventing the essential porcupine (PORCN) mediated palmitoleoylation of Wnt ligands and so generally inhibits Wnt signalling.^36^ The inhibitor XAV939 (XAV), however, targets the tankyrase (TNKS) enzyme’s ability to sequester AXIN1/2 (axis inhibition protein) in the β-catenin destruction complex, and hence specifically reduces β-catenin levels and, hence, inhibits canonical Wnt signalling.^37^ During nanovibration, treatment of cells with LGK significantly altered gene expression regulation, as illustrated in Figure 4a. Decreased mRNA expression of proinflammatory cytokine pathway genes (*BCL3*, *IL1B* and *IL6*) was seen. While β-catenin (*CTNNB1*, canonical Wnt signalling) expression was unchanged, expression of distinct non-canonical Wnts *WNT5A* decreased and expression of *WNT5B* increased.^38^ However, the non-canonical signal mediators *DVL2* and *ROR2* (tyrosine-protein kinase transmembrane receptor 2, a non-canonical Wnt receptor) were unaffected by addition of LGK. Osteogenic transcripts were also affected by addition of LGK to nanovibrationally stimulated ADSC cultures. *RUNX2* was seen to be upregulated, *SPARC* (osteocalcin, a bone ECM protein) was unchanged and *SPP1* (osteopontin, a bone ECM protein) was downregulated with the inhibitor. Osteoprotegerin, *TNFRSF11B*, expression was increased with LGK inhibition. Additional genes tested but not significantly altered following LGK treatment are shown in supplemental Figure 4. Treatment of ADSCs with XAV during nanovibration also altered gene expression (Figure 4a). While LGK decreased pro-inflammatory gene expression, treatment with XAV increased expression of *BCL3*, *IL1B* and *IL6*. XAV also increased expression of all non-canonical Wnt mediators tested (*WNT5A*, *WNT5B* and DVL2) except for *ROR2* where expression decreased; β-catenin was unaffected. For osteogenesis, *RUNX2* and *SPARC* were down-regulated by XAV inhibition with nanovibration, while *SPP1* was up-regulated. *TNFRSF11B* was strongly repressed with XAV treatment.

**Figure 4.**
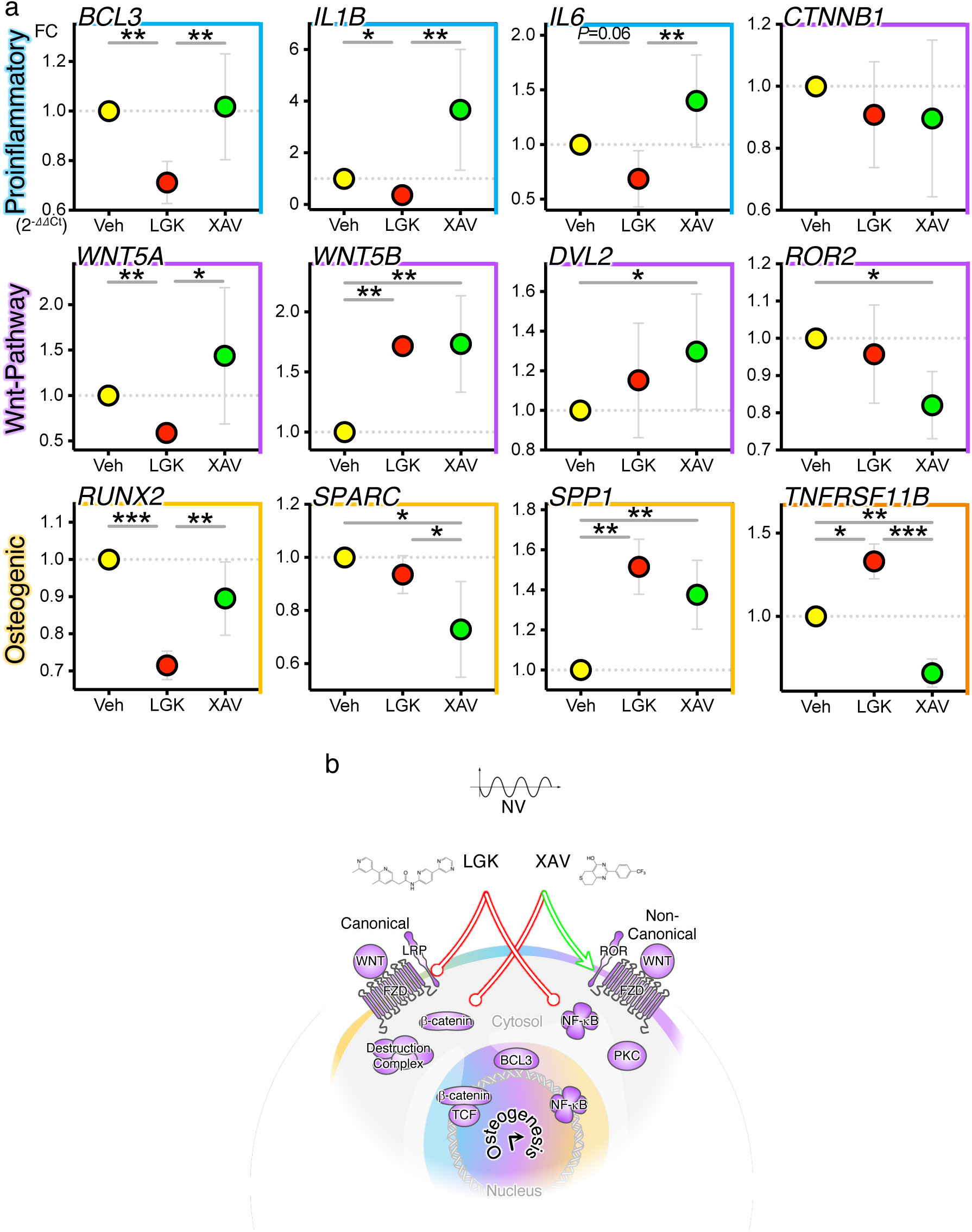
Wnt pathway inhibition shows nanovibration (NV) and non-canonical Wnt expression are synergistic. Treatment of human adipose derived stem cells (ADSCs) with Wnt pathway inhibitors LGK974 (LGK), XAV939 (XAV) or DMSO vehicle (Veh) control in the presence of nanovibration (NV), at 24 h, diverges gene mRNA expression (relative quantitative-PCR) within proinflammatory (blue), Wnt-pathway (violet), osteogenic (yellow) and anti-osteoclastogenic (orange) gene-sets (a). Repeat-measure one-way analysis of variance with Fisher’s least significant difference test of negative delta-delta Ct values (a), n = 3 independent biological replicates. Schematic representation summarising the impact of nanovibration and inhibitors affecting gene expression.

Overall, the inhibitors LGK and XAV were found to affect ADSC gene expression inversely relative to each other. It is important to note that both inhibitors, in the presence of nanovibration, did not significantly reduce β-catenin (*CTNNB1*) mRNA levels but both inhibitors *did* convergently increase *WNT5B* and osteopontin (*SPP1*) levels – a shared β-catenin-independent, non-canonical Wnt signature (Figure 4a). Additional genes less sg *BCL3* expression was seen to be unchanged by XAV, while being reduced by LGK treatment. This implies that the β-catenin-independent Wnt non-canonical pathway is involved in regulating its expression (Figure 4a). Aside from its direct role in regulating NF-κB, BCL3 activity couples with upregulated β-catenin activity in cancer and stem cells.^39–42^ Further, within osteoclasts, BCL3 can act to restrict differentiation through the RANK (Receptor activator of NF-κB)/TRAF6 (TNF receptor associated factors)/NF-κB pathway.^43^ BCL3 is also described as *the* major regulator of the senescence-associated secretory phenotype characterised by proinflammatory cytokines and matrix modifying protein secretion.^44^ This coordination of NF-κB activity by BCL3 extends to the Wnt pathway.

LGK treatment demonstrated the response to a reduction of upstream/outside-in Wnt ligand signal for both canonical *and* non-canonical branches. XAV treatment dampened downstream/cell-intrinsic canonical Wnt β-catenin-dependent activity and hence exhibited an overt non-canonical Wnt pathway signature (Figure 4b). These results are in congruence with previous studies demonstrating that LGK inhibits osteogenesis and bone formation, whereas XAV enhances them.^45–48^

### Wnt agonists further implicate non-canonical Wnt in nanovibrational ADSC osteogenesis

To further establish whether nanovibrational osteogenesis is resultant of canonical or non-canonical Wnt stimulation, ADSCs were treated with small-molecule activators in the presence of nanovibration for 24 hours (Figure 5). AMBMP hydrochloride (AMB) is a non-canonical Wnt agonist, activating Ca^2+^/calmodulin-dependent protein kinase II (CaMKII) signalling activator, regulating tubulin polymerisation, cell polarity, ciliogenesis and cell migration – characteristic of the non-canonical Wnt pathway.^49–51^ CHIR99021 (CHI), on the other hand, is an inhibitor of GSK3B (glycogen synthase kinase-3 β), itself a β-catenin inhibitor, and therefore CHI is an activator of the canonical Wnt pathway.^52^ Cells treated with AMB in the presence of nanovibration (Figure 5a) had significantly elevated expression of proinflammatory pathway genes (*BCL3*, *IL1B*, *IL6*, and *TNFA*), reduced expression of canonical Wnt components such as *CTNNB1* (β-catenin) and *WNT2* but increased expression of non-canonical Wnt components such as *DVL2* and *RSPO3* (R-spondin-3).^53^

**Figure 5.**
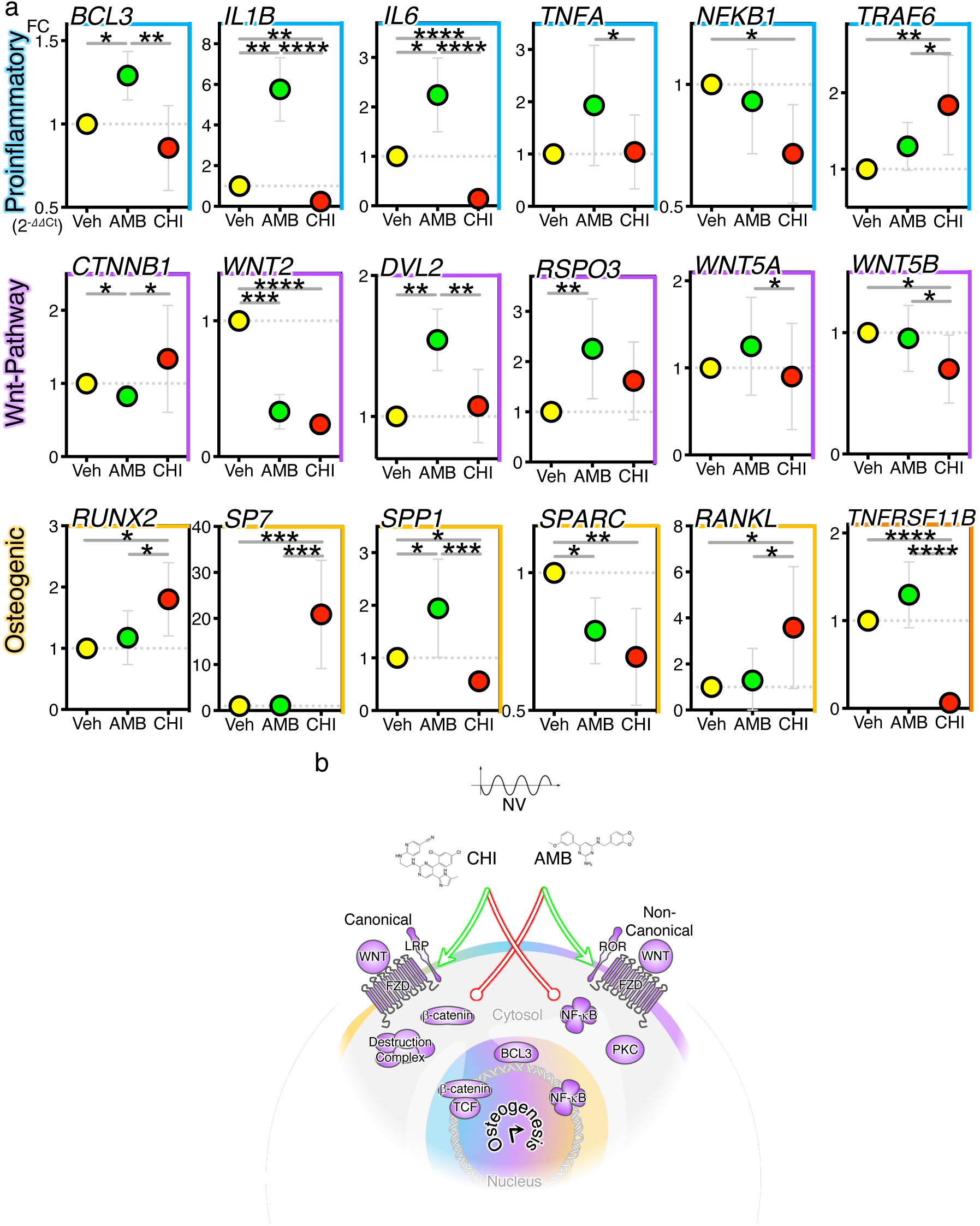
Activation of non-canonical Wnt is resembles nanovibration (NV), while canonical Wnt activation has an inverse signature. Treatment of adipose derived stem cells (ADSCs) with activators of Wnt pathway AMBMP (AMB), CHIR99201 (CHI) or vehicle (Veh) DMSO control in the presence of NV, at 24 h, diverges gene mRNA expression, assessed via relative quantitative-PCR, within proinflammatory (blue), Wnt-pathway (violet), osteogenic (yellow) and anti-osteoclastogenic (orange) gene-sets (a). Repeat-measure one-way analysis of variance with Fisher’s least significant difference test of negative delta-delta Ct values (a), n = 3 independent biological replicates. Schematic summary of results highlighting the effects of activating canonical Wnt by CHI or non-canonical Wnt by AMB and the inherent inhibition of converse pathways, in combination with NV (b).

In contrast to AMB, treatment of nanovibrated cells with CHI significantly reduced proinflammatory *IL1B*, *IL6* and *NFKB1* gene expression, increased *CTNNB1* (canonical) and decreased *WNT2* (canonical) and *WNT5B* (non-canonical) expression from the Wnt pathways. Looking at osteogenesis genes, changes included increased *RUNX2*, *SP7* (osterix, the late master osteogenic/chondrogenic transcription factor), *RANKL* (produced by mature osteoblasts to drive osteoclast formation) and decreased *SPP1 SPARC and TNFRSF11B* (the anti-osteoclastogenic cytokine osteoprotegerin) expression (Figure 5a).^35^

Thus, in the presence of nanovibration, non-canonical Wnt activation broadly enhances proinflammatory gene expression, maintains a non-canonical Wnt signature, and maintains osteogenesis (Figure 5b). On the other hand, canonical Wnt activation broadly reduces inflammatory genes, increases transcriptional drivers of osteogenesis (*RUNX2* and *SP7*) and changes the ratio of *TNFRSF11B* (reduced) to *RANKL* (increased) in favour of osteoclast formation and bone resorption. The chondrogenic SP7 is highly upregulated while the RANKL/OPG ratio is indicative of osteoclast recruitment.

It is, therefore, seen that CHI induces large changes in ADSC-osteogenic phenotype while AMB treatment results in little change in this phenotype over and above nanovibration. This suggests that further stimulation of non-canonical Wnt has little effect in line with our hypothesis that nanovibration already induces non-canonical Wnt. Stimulation of canonical Wnt, however, drives further changes to ADSC phenotype.

### Bcl3 deletion perturbs the Wnt and inflammatory genes and increases osteogenesis

The observation that nanovibrational stimulation specifically induces BCL3 expression which in turn influences NF-κB related inflammatory expression and the osteogenic phenotype through non-canonical Wnt (demonstrated by pan-Wnt inhibition using LGK to suppress this pathway, while targeting canonical Wnt with XAV leads to no suppression, Figures 3 & 4), lead us to identify BCL3 as potentially orchestrating these pathways and as a target for functional knockout and overexpression. We decided to study the impact of BCL3 knockout on MM induced osteogenesis using *Bcl3^-/-^* osteoblasts from mouse calvaria compared to wild type (WT) cells (Figure 6a). *Bcl3* knockout led to a large number of differentially expressed genes as measured by RNA-seq (Figure 6a). Crucially, gene ontology analysis of upregulated genes (Figure 6b) revealed prostaglandin (PG) receptor activity to be most upregulated, with Wnt-binding also among the most enriched gene-sets. Indeed, the prostaglandin pathway is associated with the Wnt pathway and BCL3 is a key determinant of this pathway.^40^ Other enriched gene-sets were seen to be involved in matrix function and signalling, including heparin sulphate proteoglycans (HSPG), fibronectin (FN), metalloexopeptidase activity (MXP), heparin binding (Hep), transmembrane receptor protein tyrosine kinase activity (RTK) and growth factor activity (GF), and in preventing damage from oxidative stress, such as glutathione peroxidase activity (GSHP). Looking at individual genes (Figure 6c), significant upregulation of proinflammatory, Wnt-pathway and osteogenic genes were identified with osteogenic differentiation in *Bcl3^-/-^*, compared to WT osteoblasts. This helps to place BCL3 as a regulator of Wnt and osteogenesis, alongside its role as an inhibitor of NF-κB activity.^30^ At the transcript level, BCL3 depletion appears to recapitulate the non-canonical Wnt signature observed with NV and inhibition by XAV (Figure 6c,f).

**Figure 6.**
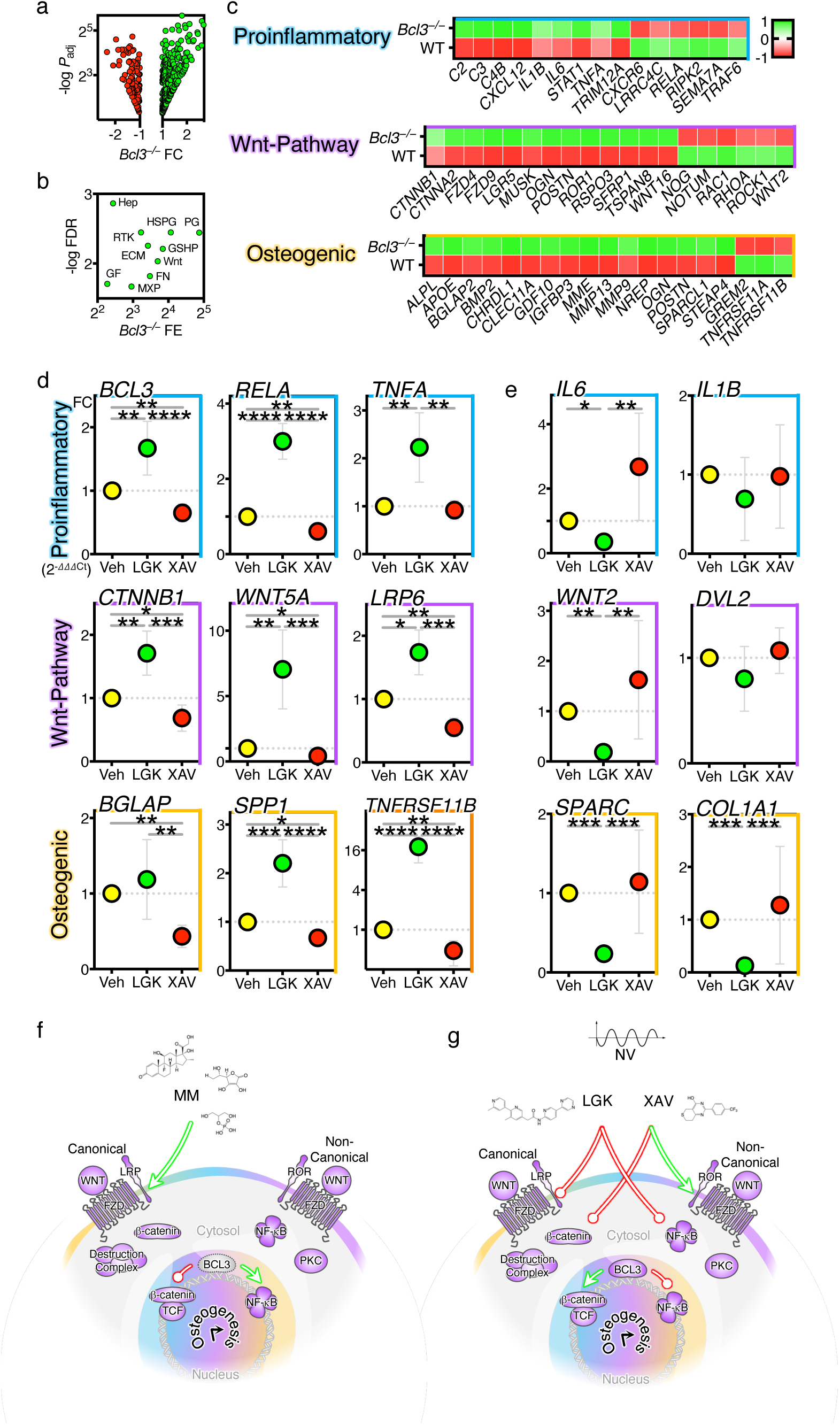
BCL3 is at the nexus of the proinflammatory, Wnt pathway and osteogenic gene-sets and coordinates the switch between canonical and non-canonical Wnt signalling. BCL3-deleted (Bcl3^-/-^) and WT murine calvarial osteoblasts were stimulated with MM for 24 h and their gene expression transcripts were quantified via RNA-sequencing (GSE125153). Volcano plot of Bcl3^-/-^ fold-change (FC) compared to WT and adjusted P values of differential expression (a), gene ontology Bcl3^-/-^ fold-enrichment (FE) compared to WT and -log false discovery rate (FDR) value (b), and expression Z-score heatmaps for selected gene-sets (c), n = 3 independent biological replicates. Ontology abbreviations: prostaglandin receptor activity (PG), heparan sulphate proteoglycan binding (HSPG), glutathione peroxidase activity (GSHP), Wnt-protein binding (Wnt) fibronectin binding (FN), extracellular matrix structural constituent conferring tensile strength (ECM), transmembrane receptor protein tyrosine kinase activity (RTK), metalloexopeptidase activity (MXP), heparin binding (Hep), and growth factor activity (GF). Human ADSCs were supplemented with BCL3-mimetic peptide, alongside Wnt inhibitors LGK974 (LGK) or XAV939 (XAV), in the presence of nanovibration for 7 days and transcript expression accounted for baseline no-peptide controls (d and e). Repeat-measure one-way analysis of variance with Fisher’s least significant difference test of negative delta-delta-delta Ct values, n = 4 independent biological replicates. Schematics highlighting the net effects of BCL3 deletion in stromal cells in the presence of metabolite media (MM) stimulation (f), and BCL3 overexpression in nanovibration (NV) stimulation and in the presence of either LGK or XAV (g).

To further explore the role of BCL3, observed to be differentially expressed during nanovibration and Wnt inhibition, a BCL3 mimetic peptide,^26^ was supplemented to ADSCs for 7 days with nanovibration-stimulated osteogenesis (Figure 6d,e). It was seen that across inflammatory drivers (*BCL3*, *RELA* and *TNFA*), cytokines (*IL6* and *IL1B*), canonical and non-canonical Wnt-related drivers (*CTNNB1*, *WNT2*, *WNT5A*, *LRP6* and *DVL2*), and osteoblast-related transcripts (*TNFRSF11B*, *SPP1*, *BGLAP*, *SPARC*, *COL1A*), there was large dysregulation with the use of the pan-Wnt inhibitor LGK with BCL3-overexpressing nanovibrated ADSCs; this tended towards up-regulation of genes in BCL3 deletion with MM (Figure 6d) and in BCL3 overexpression with NV (Figure 6e). At the same time, there was very little change observed with use of the canonical-Wnt inhibitor XAV in the same cultures (Figure 6d,e). Additional genes tested but not significantly changed are shown in supplemental Figure 5. This more clearly links BCL3 and non-canonical Wnt in osteogenic cultures, here driven by nanovibrational stimulation (Figure 6g).

### Nanovibration regulates osteoclastogenic genes in a BCL3-dependent manner

Throughout this study, we have seen changes in regulation of osteoprotegerin (*TNFRSF11B*), which is up-regulated during nanovibrational osteogenesis and down-regulated with pan-Wnt inhibition (but not with canonical-Wnt inhibition) and not affected by non-canonical-Wnt agonism. As noted, osteoprotegerin is secreted by maturing osteoblasts to prevent osteoclast formation and bone resorption. Together, this suggests that expression of osteoprotegerin is stimulated by the non-canonical-Wnt pathway during nanovibrational stimulation and that further non-canonical-Wnt stimulation does not produce more osteoprotegerin expression. Previously, it has been reported that similar nanovibrational stimulation prevents human osteoclast formation and bone resorption^54^, and this agrees with our osteoprotegerin data.

As we develop our hypothesis that non-canonical Wnt, regulated by BCL3, is important in nanovibrational osteogenesis, we asked if osteoclast formation is also affected by BCL3. To study this, murine bone marrow derived CD14^+^ monocytes from WT and *Bcl3^-/-^*mice were primed to undergo osteoclastogenesis, with or without nanovibration (Figure 7). Cells were cultured up to 5 days with osteoclastogenic cytokines M-CSF and RANKL to permit macrophage fusion into osteoclasts.^55^ Nanovibration increased gene expression at day 3 of select osteoclastogenic functional genes, namely, *OSCAR* (osteoclast-associated immunoglobulin-like receptor) and *OCSTAMP* (osteoclast stimulatory transmembrane protein) in WT cells, but not in *Bcl3^-/-^* cells (Figure 7a), as well as in other marker genes (Supplemental Figure 6). Thus, nanovibration was seen to increase osteoclast-related transcripts, as was previously observed.^54^ *Bcl3* deletion reduced the expression of osteoclastic genes during nanovibration, identifying BCL3 as a mechanoresponsive protein in osteoclasts.

**Figure 7.**
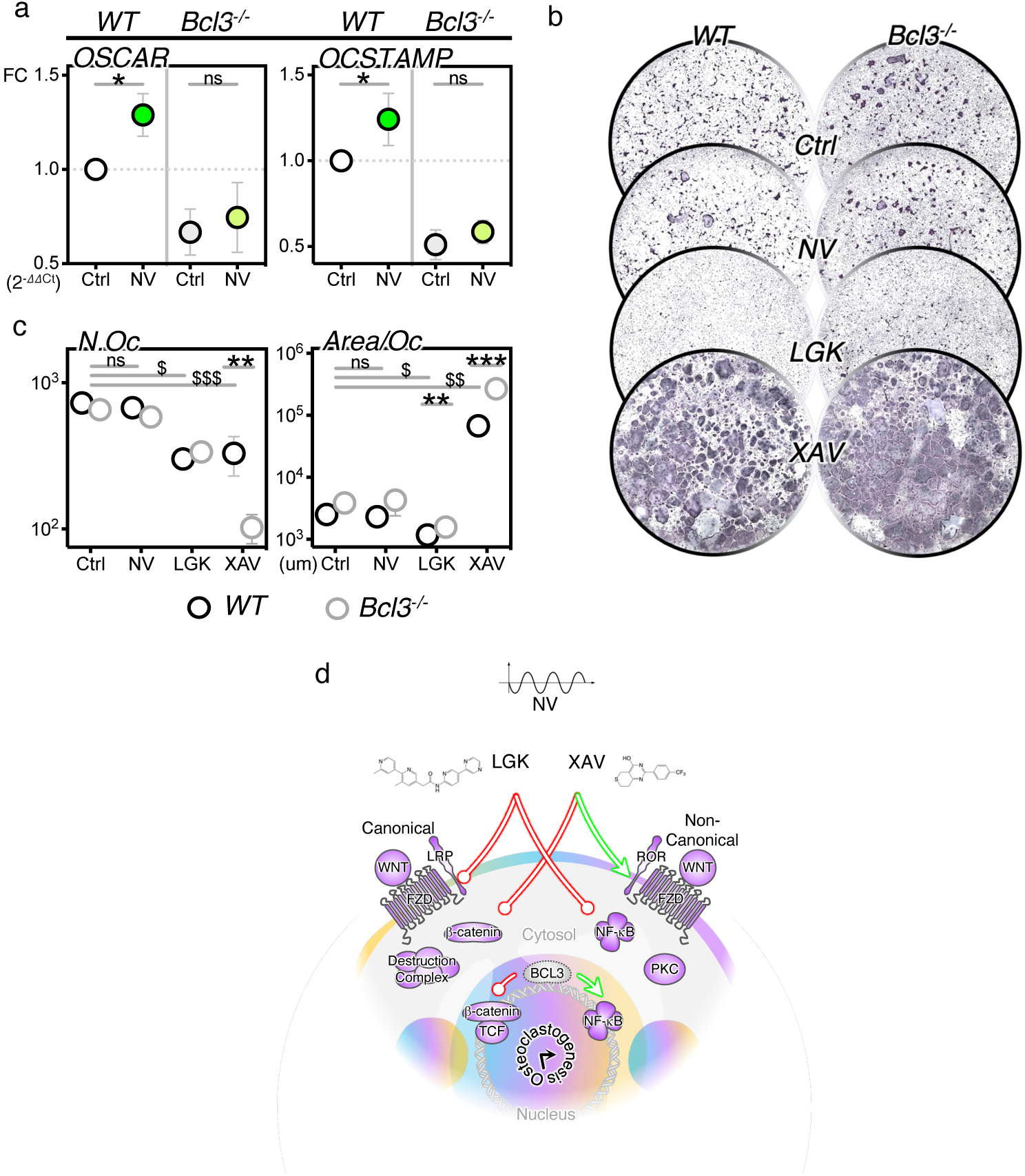
In osteoclasts, BCL3 absence is also synergistic to the non-canonical Wnt pathway. Osteoclastogenesis of BCL3-deleted (Bcl3^-/-^) and WT utilised murine CD14^+^ monocyte precursor cells supplemented with M-CSF and RANKL cytokines, for 5 days, with or without nanovibration (a), LGK974 (LGK) or XAV939 (XAV) (b,c). Osteoclast marker gene expression of WT and Bcl3^-/-^ cells, in presence or absence of nanovibration (NV) determined via relative quantitative-PCR (a). Repeat-measure one-way analysis of variance with Fisher’s least significant difference, n = 4 independent biological replicates (a). Purple TRAP (tartrate-resistant acid phosphatase) stained osteoclast cells imaged under brightfield, with circular image diameter = 32 mm, counted and area per cell quantified (b). Repeat-measure one-way analysis of variance with Fisher’s least significant difference test across conditions ($) and Student’s t-test between genotypes (*), n = 4 independent biological replicates (c).

As functional osteoclast differentiation and activity is correlated to their number and size (from macrophage fusion), TRAP (tartrate-resistant acid phosphatase, an osteoclast activity marker) staining of WT and *Bcl3^-/-^* cells, treated either with nanovibration, with the Wnt inhibitors LGK or XAV, or as untreated controls were assessed (Figure 7b,c). For WT cells, as in previous work,^54^ increases in osteoclast gene expression did not translate to more or larger osteoclasts (Figure 7c). While in human cells, there was a significant decrease in osteoclast (fused cells containing three or more nuclei) number and size post vibration, with murine cells, this was reduced to a trend (Figure 7c). There appeared to be no difference between control and nanovibrational conditions with *Bcl3^-/-^* cells (Figure 7c).

The pan-Wnt inhibitor, LGK, reduced osteoclast formation equally in WT and *Bcl3^-/-^* (Figure 7b,c). The canonical-Wnt inhibitor XAV did not affect osteoclast number relative to LGK, however it increased osteoclast size with WT cells, and in *Bcl3*^-/-^ cells increased osteoclast size to the point of decreasing individual cell number (Figure 7c). This suggests that canonical Wnt activity is associated with BCL3 presence. Inhibition of canonical Wnt by itself drives up osteoclast formation, indicating that canonical Wnt can help reduce osteoclast formation in an osteo-permissive environment, such as nanovibration provides. BCL3 inhibition synergises with canonical Wnt inhibition, exhibiting massive increases in cell size and suggesting increased osteoclast fusion and enhanced resorptive phenotype. This synergy indicates that BCL3 is also involved in an osteoclast-repressive mechanism, alongside nanovibration.

Enhanced osteoclastogenesis by XAV has previously been shown to coincide with β-catenin sequestration.^45^ In support of these results, *BCL3*, β-catenin (*CTNNB1*), *IL6*, *IL1B*, *RELA* and the Wnt transcription factor *TCF7L2* were recently identified as a major co-expressing gene cluster especially upregulated at 4 hours to 1 day but then dampening expression with increasing differentiation of human osteoclasts.^56^ This gene-set was also associated with the most estimated bone mineral density-associated single-nucleotide polymorphisms (SNPs) and was most expressed in early fracture response.^56^ Thus, XAV’s inhibition of the canonical Wnt/β-catenin pathway is likely to increase osteoclastogenesis, as we see here.

Roles for BCL3 in regulating non-canonical Wnt are less clear as there is a trend of reduced osteoclast formation with/without BCL3 deletion. However, with the tools we have, it is hard to draw any conclusion beyond postulating canonical and non-canonical Wnt might be in competition, as we used a pan-Wnt inhibitor to infer changes from the canonical-Wnt inhibitor and we saw a mild, opposite, change.

## Conclusion

Herein we have we elucidated the emergence of a nanovibration-related molecular signalling axis incorporating proinflammatory NF-κB and non-canonical Wnt activation with positive osteogenic differentiation. We establish BCL3 as a central player in regulation of inflammation, non-canonical Wnt and osteogenesis. Until now, the nanovibrational mechanism of osteogenesis has been vague and detail has been hard to draw. Inference has been made previously to integrins and cell adhesion, cation channels, YAP translocation and Wnt.^57^ Here, we show that inhibition of non-canonical Wnt causes reduction in nonvibrational osteogenesis, while inhibition of canonical Wnt does not. Further, if BCL3 is deleted, these central pathways involved in nanovibrational osteogenesis become dysregulated. This is the first study to robustly probe the mechanism of nanovibrational mechanotransduction in bone formation. Critically, we show that nanovibrational biomechanic stimulus is sufficient to engage this novel, emergent mechanotransductive axis for osteogenic differentiation.

## Methods

### Laser Doppler Vibrometry

The vibrational response of the nanoamplitude vibrational bioreactor was measured using a scanning laser Doppler vibrometer (LDV) (Model MSA-100-3D, Polytec GmbH, Germany). All LDV scans were performed at 1 kHz, with a ±1.4 V sinusoidal signal generated by the LDV’s internal signal generator and then amplified (40X) by a Behringer KM-750 amplifier (Behringer, Germany) drive the bioreactor top plate. All LDV scans were conducted with the following settings: sampling frequency, 6.25 MHz; bandwidth, 5 kHz; fast Fourier transform lines, 3200; resolution, 1.56 Hz; and averaging of 10 measurements per point. During cell culture, the nanovibrational bioreactor is powered by its own power supply unit.^14^

A 6-well plate and 4-well chamber slide were separately magnetically attached, using self-adhesive rubber magnet, to the bioreactor’s top plate for these measurements (Figure 1a and supplemental Figure 1a). For the 6-well plate setup, an identical circular measurement grid of 41 points was created and centred across the bottom (cell culture surface) of each well in PSV software (Version 8.51, Polytec GmbH, Germany), this was done to measure the instantaneous displacement (Figure 1b) and maximum displacement (Figures 1c and 1d) at 1 kHz. For the 4-well chamber slide, an identical rectangular grid of 45 equally spaced points was created for each well. The LDV’s field of view was too small to cover the entire 6-well plate, so only 4 wells are depicted in Figure 1b. However, all 6 wells are shown in Figure 1c, as an R script (version 4.2.3) was created to examine the displacement pattern across each well in detail, enabling two separate scans to be stitched together.

To determine the displacement and phase difference between the bottom of the well and the cell media’s surface, a smaller circular measurement grid of 13 points per well was created and centred across the bottom of each well. These same 13 points were used to take measurements from the liquid’s surface. The smaller measurement grid was used to avoid taking measurements from the meniscus. The displacement and phase differences were then determined by subtracting the liquid’s surface measurement from the corresponding well-bottom measurement.

The longitudinal pressure change caused by the 1 kHz, 30 nm amplitude vibration is visualised for two complete wavelengths in Figure 1h. In this plot, the time-varying displacement is depicted by the solid black line, and the acoustic wave pressure distribution is plotted by a point-cloud representation, where dot density and colour represent the spatial pressure variation. Further, the zoomed-in region depicts the longitudinal pressure change through the cell media (height 5 mm). This pressure change was calculated using:

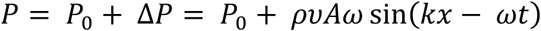

Where 𝑃 is total pressure, 𝑃_0_ ambient pressure (100 kPa), Δ𝑃 the pressure variation due to nanovibration. 𝜌 is the density of cell media (1000 kg m^-3^), 𝜐 the speed of sound in cell media (1500 m s^-1^), 𝐴 the amplitude (30 nm), 𝜔 the angular frequency (2000𝜋 s^-1^), 𝑘 the wavenumber (= 2𝜋⁄𝜆), where 𝜆 is the wavelength (1.5 m), 𝑥 the position (m) and 𝑡 the time (s).

We analysed the velocity and time data directly measured by the LDV to estimate the accelerative forces acting on whole cells during vibration (Figure 1i). The accelerative force experienced by a typical adherent cell is the product of the mass of the column of cell media that it supports and the acceleration, i.e. Newton’s second law. The mass of the column was estimated by taking the average area of a stromal cell/osteoblast to be 1000 µm^2,58^ the height of the column to be 5 mm, and the density of the media to be 1000 kg m^-3^. The acceleration was acquired by differentiating the velocity and time data directly measured by the LDV.

A single-point laser interferometer (Model SP-S SIOS Meßtechnik GmbH, Germany) was used to acquire the displacement-frequency spectrum of the nanovibrational bioreactor with culture ware (Figure 1e). In this experiment, the bioreactor was powered by the in-house designed power supply unit.^14^ The frequency spectrum was acquired from the central point of each well, 10 measurements were taken per well (6-wells or 4-wells, respectively).

### Cell Culture of ADSCs

Human adipose-derived stromal/stem cells (ADSCs), isolated from excess adipose tissue from female liposuction patients, were commercially obtained as cryopreserved vials (Histocell Regenerative Medicine). In basal condition, cells were cultured in Dulbecco’s Modified Eagle’s medium (DMEM) with 4.5 g/L glucose (D5671), supplemented with 10% foetal bovine serum (FBS; F9665), 2 mM L-glutamine (25030), 500 U/mL penicillin and 20 ug/mL streptomycin (P/S) antibiotics (15140), minimal essential medium non-essential amino acids (MEM NEAA) solution (11140), 5 mM sodium pyruvate (11360) and 0.5 ug/mL amphotericin B antimycotic (15290). In certain instances, clonal expansion was accelerated using MesenPRO RS Basal Medium (12747) with Growth Supplement (12748). To preserve cell stemness and response to stimuli, cell clonal expansion of ADSCs for sufficient cell number was limited up to a maximum of 5 passages, for any given donor, and confluency did not exceed 80% in T-75, T-150 flasks. All reagents were procured from Thermo-Fisher Scientific or Merck.

### Treatment Conditions

To serum starve cells during or prior to experimental treatment that lasted up to 1 day, Opti-MEM Reduced Serum Medium (31985; Thermo-Fisher Scientific) supplemented with 2% FBS and P/S was used. Transcriptional assays were conducted in 6-well plates. Nanovibrational (NV) stimulus was delivered to adherent cells in plasticware magnetically attached to the piezoelectrically-driven bioreactor device surface, with displacement calibrated to ±30 nm oscillating amplitude at a 1 kHz frequency.^59^ Osteogenic metabolite media (MM) was comprised of 350 μM ascorbic-2-phosphate, 10 nM dexamethasone and 5 mM β-glycerophosphate. Various molecules supplementary to the above basal media were utilised as treatment conditions during the course of the study. Small molecule inhibitors LGK974, 2-(2’,3-Dimethyl-[2,4’-bipyridin]-5-yl)-N-(5-(pyrazin-2-yl)pyridin-2-yl)acetamide (CAS: 1243244-14-5); XAV939, 3,5,7,8-Tetrahydro-2-[4-(trifluoromethyl)phenyl]-4H-thiopyrano[4,3-d]pyrimidin-4-one (CAS: 284028-89-3); AMBMP, 2-Amino-4-(3,4-(methylenedioxy)benzylamino)-6-(3-methoxyphenyl)pyrimidine (CAS: 2095432-75-8); CHIR99201, 6-[[2-[[4-(2,4-Dichlorophenyl)-5-(5-methyl-1H-imidazol-2-yl)-2-pyrimidinyl]amino]ethyl]amino]-3-pyridinecarbonitrile (CAS: 252917-06-9); and Y27632, trans-4-[(1R)-1-Aminoethyl]-N-4-pyridinylcyclohexanecarboxamide (CAS: 129830-38-2) were procured from R&D Systems, dissolved in DMSO and used at a final working concentration of 10 μM. BCL3 mimetic peptide BDP2 with the protein sequence YGRKKRRQRRAAVYRILSLFKLGSR was used at a final concentration of 16 μM.^26^

### Cell Culture Osteoclast

Bone marrow from 8-week-old male WT and Bcl3^-/-^ mice^30^ was aseptically isolated and suspended into complete MEMα (22571; ThermoFisher Scientific). CD14^+^ monocytes were isolated via immunomagnetic selection using EasySep Mouse Monocyte Isolation Kit (Stemcell Technologies). Non-adherent monocytes were expanded overnight with 30 ng/mL M-CSF (416ML) and subsequently seeded into 96-well plates at 10^5^ cells/well supplemented with 50 ng/mL M-CSF and 50 ng/mL RANKL (462-TEC), both cytokines procured from R&D Systems, and further cultured for 4 days to observe mature osteoclastogenesis. Tartrate-resistant acid phosphatase (TRAP) enzymatic stain was performed, in accordance with Leukocyte Acid Phosphatase Kit (387A) from Merck. Assessment of bona fide osteoclasts and area calculations was conducted with Fiji.

### Immunoblotting

Immunoblotting was performed to evaluate the effect of nanovibrational stimulation on Rho-associated coiled-coil kinases. The experiment comprised of one control and two treatment groups. Control group did not receive nanovibrational stimulation. Both treatment groups were subjected to nanovibrational stimulation. One of the treatments was subjected to ROCK-inhibitor Y27632, trans-4-[(1R)-1-Aminoethyl]-N-4-pyridinylcyclohexanecarboxamide dihydrochloride (CAS: 331752-47-7), procured from R&D Systems, whereas the other was not. ADSCs from three different donors were seeded at a density of 2000 cells/cm^2^ in T-75 flasks. Both treatment groups were subjected to nanovibrational stimulation for 7 days. One day prior to experimental endpoint, complete media from control and both treatment groups were aspirated, cells were washed with DPBS and opti-MEM reduced serum media was added overnight to sync cell cycles. The following day, Y27632 was added to one of the treatment groups at 10 μM concentration for 4 hours. Cells in all experimental groups were then washed with ice-cold PBS and lysed using Radioimmunoprecipitation assay (RIPA; 89900) buffer containing Halt protease and phosphatase inhibitors (78438) at final concentration of 1X, procured from ThermoFisher Scientific. Proteins were extracted by centrifugation at 13,000 RPM for 30 minutes at 4°C. Total proteins were quantified using Bicinchoninic assay (BCA; 23225) and equal amounts of proteins were loaded on Bolt Bis-Tris 4-12% polyacrylamide gels (NW04122) containing reducing agent (NP0004) and sample buffer (NP0007) at 1X final concentration, all procured from ThermoFisher Scientific. Proteins were then transferred to PVDF membrane at 25V and 160 mA for 70 minutes in a cooled environment and blocked using 5% non-fat dry milk (NFDM) or bovine serum albumin for one hour at room temperature. Membranes were incubated with protein-specific antibodies for overnight at 4°C. The phospho-Cofilin (Ser3) polyclonal antibody (29715-1-AP) and the CDC42 polyclonal antibody (10155-1-AP) were both procured from Proteintech, and the phospho-Myosin Light Chain 2 (Ser19) polyclonal antibody (3671) was procured from Cell Signaling Technology. The following day HRP-conjugated secondary antibodies diluted in 5% NFDM and incubated with membranes for one hour at room temperature. Proteins were then imaged using PicoPlus chemiluminescent substrate (34578).

### Immunofluorescence

Immunofluorescence was performed to study the effect of nanovibrational stimulation on cell cytoskeleton by activating ROCK signalling. Cells were seeded at a density of 2000 cells/cm^2^ in LabTek chambered slides. Experimental groups comprised of a control group without nanovibrational stimulation, and two treatment groups subjected to nanovibrational stimulation. One of the treatment groups was subjected to ROCK inhibition using Y27632, while the other groups was not. ROCK inhibition was performed following the above-mentioned procedure. After 7 days, cells were washed with warm DPBS and fixed using 4% formaldehyde for 10 minutes at room temperature. Cells were permeabilised using 0.2% TritonX-100 and then blocked for one hour at room temperature using 2% BSA. Cells were incubated with phosphorylated myosin light chain 2 antibodies overnight at 4°C. The following day, cells were incubated with Alexa Fluor 647 secondary antibodies and Alexa Fluor 568 Phalloidin to immunostain filamentous actin (F-Actin) for visualising stress filaments. Cells were mounted on a glass slide using Prolong Gold mounting media containing DAPI, dried overnight at room temperature followed by imaging the following day with a Zeiss LSM 880. Samples were washed three times for 5 minutes each using DPBS after fixation, permeabilization, primary and secondary antibody incubations.

### Animals

*Bcl3*^-/-^ and WT littermates were bred under standard conditions, and all animal studies received ethical approval from the University of Glasgow and were performed under a UK Home Office license.

### Quantitative PCR

Cells were lysed with RLT buffer supplemented with 1% β-mercaptoethanol, RNA was purified with the RNeasy Mini Kit (74104), cDNA synthesis was performed with the QuantiTect Reverse Transcription Kit (205311), quantitative reverse transcription polymerase chain reaction, qRT-PCR, was performed with the QuantiTect SYBR Green PCR Kit (204143) – all procured from Qiagen. Relative quantitation was performed with normalisation to ribosomal protein L13a (*RPL13A*) as the constitutively expressed gene in ADSCs and glyceraldehyde-3-phosphate dehydrogenase (*GAPDH*) in osteoclasts.

**Table 1.**
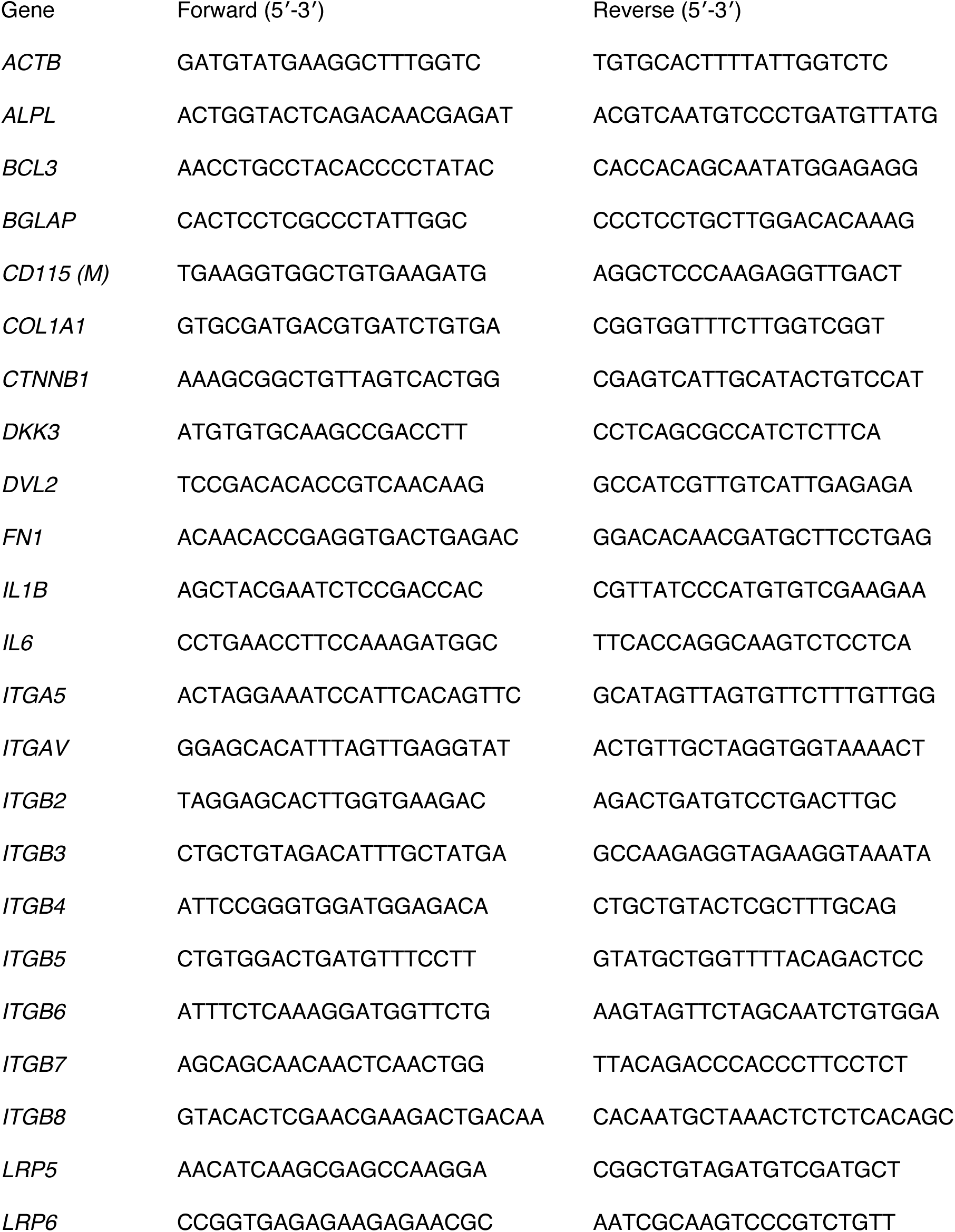
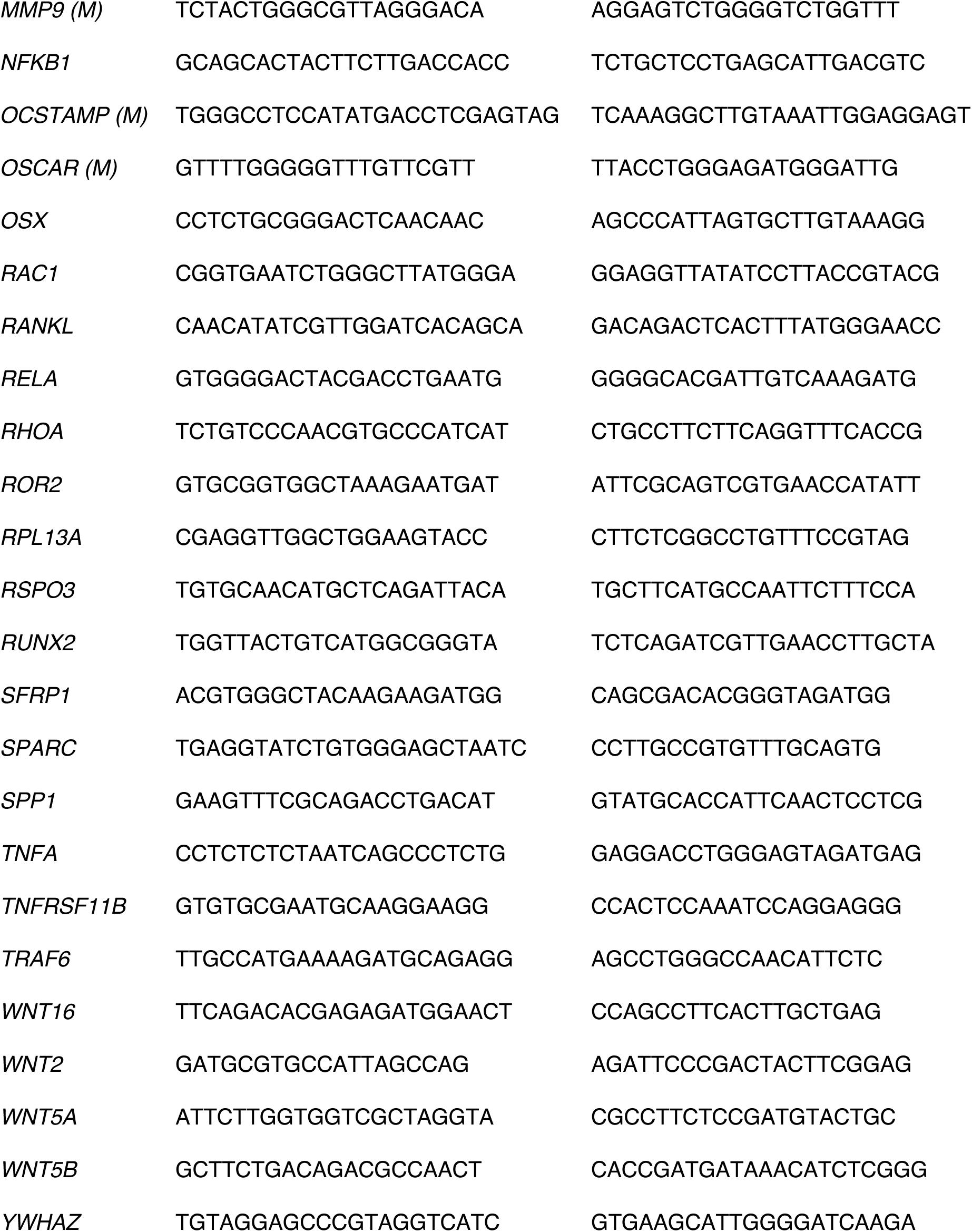
List of primers used for qPCR reactions. Primers demarcated ‘(M)’ are specific to murine cells and all other primers are specific for human cells.

### RNA-seq

RNA-sequencing data (Gene Expression Omnibus accession GSE125153) was experimentally produced by isolating murine calvarial osteoblasts from WT and Bcl3^-/-^ mice (n = 3) and stimulating differentiation with 50 μg/mL L-ascorbic acid and 2 mM β-glycerophosphate, for 24 hours. The SMART-seq v4 (634891) and Library Prep Kits from Clontech were utilised. Sequencing was performed at 40 million, 75 bp paired-end reads per sample via HiSeq 4000 (Illumina) Cufflinks and DESeq2 was used to calculate differential expression and adjusted P value of <0.05 was considered significant.^30^

### Statistical Approach

Statistical analyses utilised GraphPad Prism 10. Data was tested for normality using the D’Augustino-Pearson omnibus normality test. All tests were two-tailed. In data where the sample size (n) was insufficient for a normality test, normality was assumed. In two-group data where the n was insufficient for a normality test, the Student’s t test with the assumption of an identical standard deviation (SD) was used. For normally distributed data with two groups, the unpaired t test with Welch’s correction or the paired ratio t test was applied. For two-group data failing the normality test, the unpaired Mann-Whitney test was used. Data with multiple groups and a single variable were compared using analysis of variance (ANOVA), followed by multiple comparisons using Fisher’s least significant difference test. P-values <0.05 were considered significant, throughout.

## Acknowledgements

Healikick Project Consortium. This project has received funding from the European Union’s Horizon 2020 research and innovation programme under grant agreement No 874889.

## Notes

### Competing Interest Statement

The authors have declared no competing interest.

