## Supplemental Figures for "Nanovibrational stimulation of osteogenesis engages non-canonical Wnt signalling and NF-κB regulator BCL3 as a mechanotransducer"

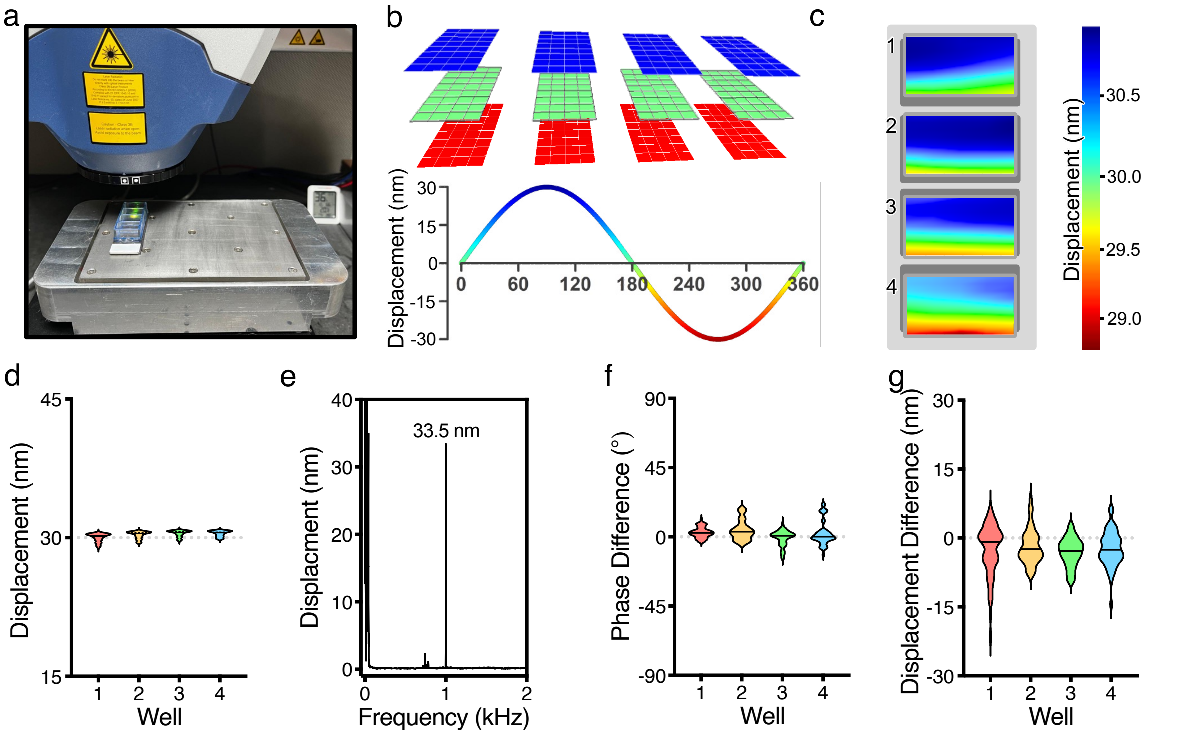


***Supplemental Figure 1** | Scanning laser Doppler vibrometry-based characterisation of the nanovibrational bioreactor. The nanovibrational bioreactor and 4-chamber slide magnetically coupled to the top plate (a). The instantaneous z-displacement at three separate points (0 (green), 90° (blue) and 270° (red) of one vibration cycle, measured at 1 kHz (b). Colour variance map of maximum z-displacement (90° of vibration cycle) in relation to position within wells of culture plate at 1 kHz (c). Violin plot of the displacement at 1 kHz for each slide chamber, with 45 measurement sites per chamber (d). The average frequency spectrum of the z-displacement from 0.4 to 3000 Hz from all*

*measurement points, with a primary peak at 1 kHz (e). The phase difference (f) and displacement difference (g) between the bottom of each chamber and the cell media's surface, showing variation.*

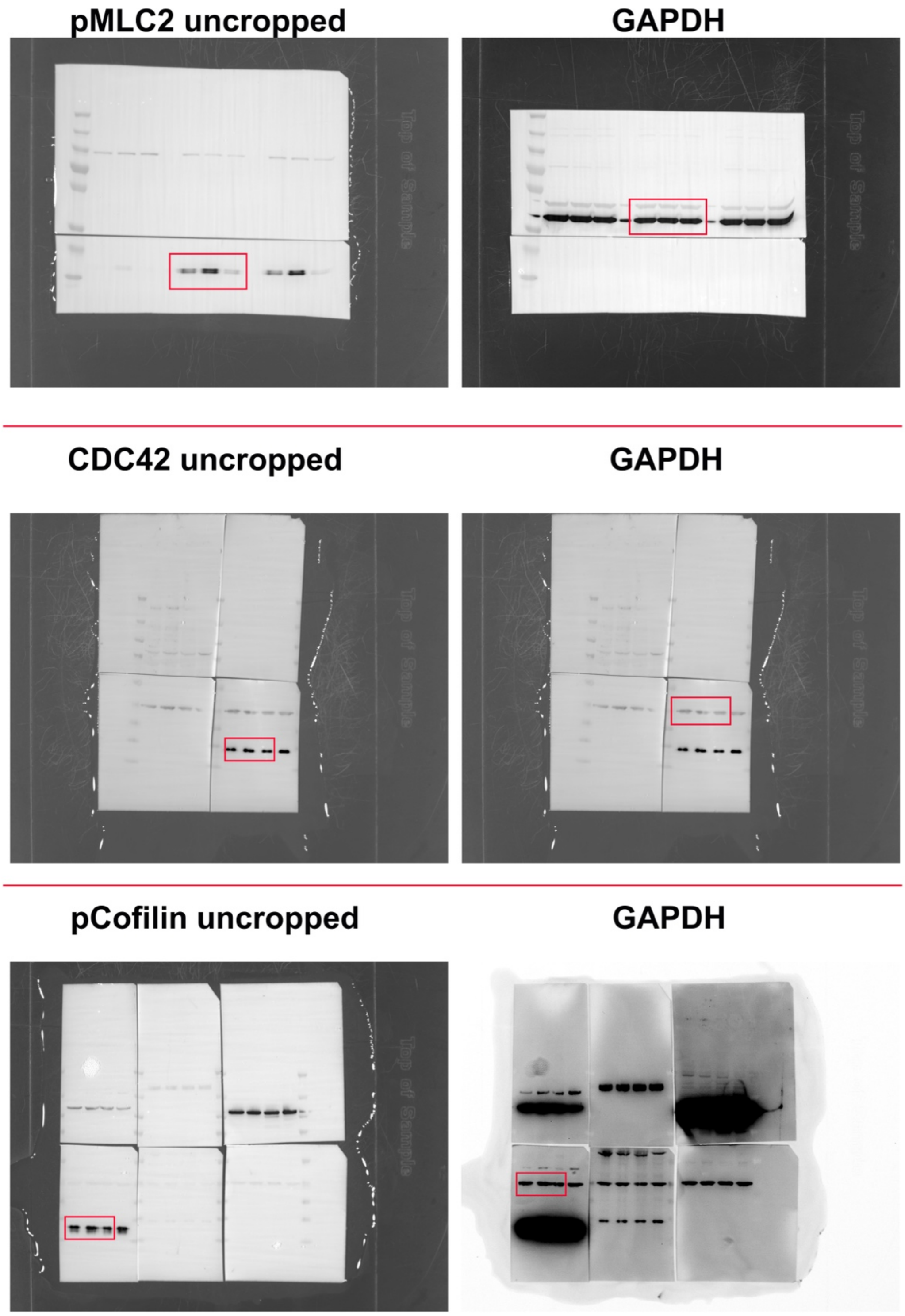

***Supplemental Figure 2** I Complete uncropped images of immunoblots of p-MLC2, CDC42and p-Cofilin, and their GAPDH counterparts, showing full context of the analysed data. Red boxes denote the images extracted for the primary Figure 2.*

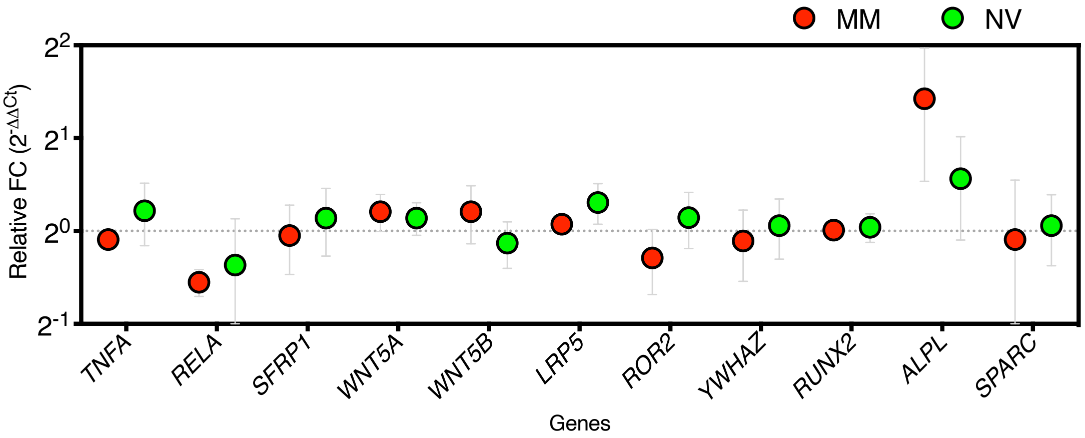


*Supplemental Figure 3 I Adipose derived stromal cells (ADSCs) that were nanovibrated (NV) compared to metabolite medium (MM), stimulated at day 7. Repeat-measure one-way analysis of variance with Fisher's least significant difference test incorporating basal unstimulated group of negative delta-delta Ct values, n = 3 independent biological replicates.*

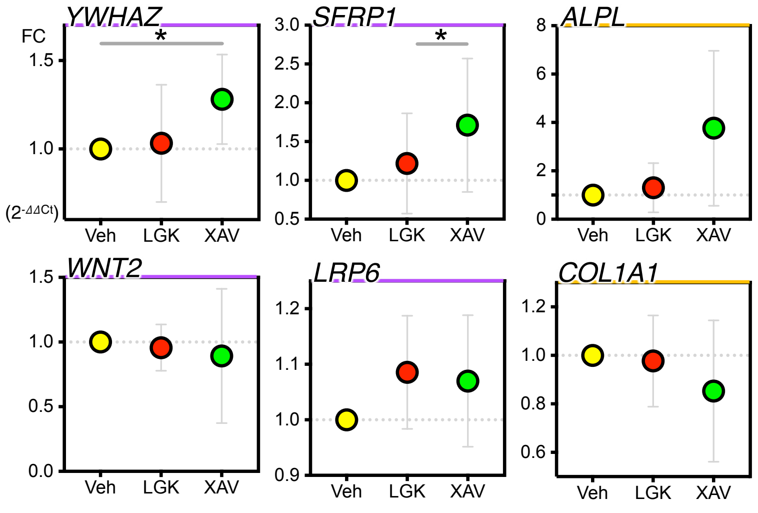


*Supplemental Figure 4 I Treatment of adipose derived stem cells (ADSCs) with Wnt pathway inhibitors LGK974 (LGK), XAV939 (XAV) or vehicle (Veh) DMSO control in the presence of*

*nanovibration (NV), at 24 h. Repeat-measure one-way analysis of variance with Fisher's least significant difference test of negative delta-delta Ct values, n = 3 independent biological replicates.*

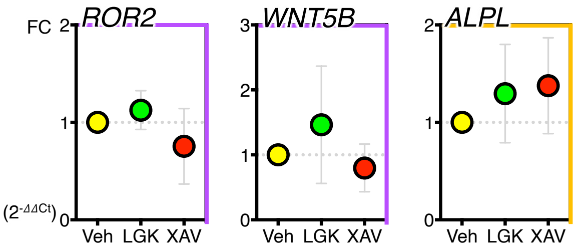


*Supplemental Figure 5 I ADSCs were cultured with BCL3-mimetic peptide (BCL3 overexpression), alongside Wnt inhibitors LGK974 (LGK) or XAV939 (XAV), in the presence of nanovibration for 7 days and transcript expression accounted for controls (d and e). Repeat-measure one-way analysis of variance with Fisher's least significant difference test of negative delta-delta-delta Ct values, n = 4 independent biological replicates.*

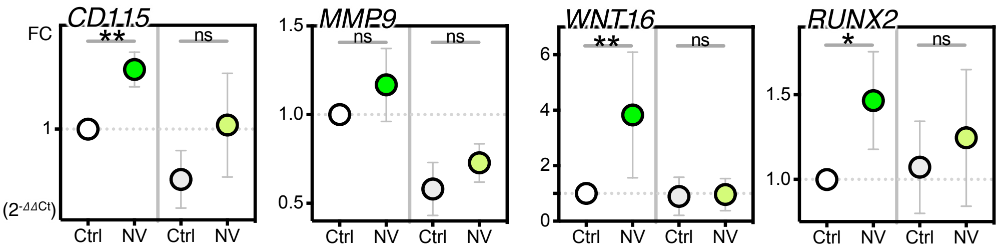


*Supplemental Figure 6 I Osteoclast marker gene expression of WT and $BCL3^{-/-}$ cells, in presence or absence of nanovibration (NV). Repeat-measure one-way analysis of variance with Fisher's least significant difference, n = 4 independent biological replicates.*
